# A Python-based optimization framework for high-performance genomics

**DOI:** 10.1101/2020.10.29.361402

**Authors:** Ariya Shajii, Ibrahim Numanagić, Alexander T. Leighton, Haley Greenyer, Saman Amarasinghe, Bonnie Berger

## Abstract

Exponentially-growing next-generation sequencing data requires high-performance tools and algorithms. Nevertheless, the implementation of high-performance computational genomics software is inaccessible to many scientists because it requires extensive knowledge of low-level software optimization techniques, forcing scientists to resort to high-level software alternatives that are less efficient. Here, we introduce Seq—a Python-based optimization framework that combines the power and usability of high-level languages like Python with the performance of low-level languages like C or C++. Seq allows for shorter, simpler code, is readily usable by a novice programmer, and obtains significant performance improvements over existing languages and frameworks. We showcase and evaluate Seq by implementing seven standard, widely-used applications from all stages of the genomics analysis pipeline, including genome index construction, finding maximal exact matches, long-read alignment and haplotype phasing, and demonstrate its implementations are up to an order of magnitude faster than existing hand-optimized implementations, with just a fraction of the code. By enabling researchers of all backgrounds to easily implement high-performance analysis tools, Seq further opens the door to the democratization and scalability of computational genomics.

## Introduction

The vast growth of next-generation sequencing (NGS) data has provided us with a new understanding of many biological phenomena. As such technologies rapidly evolve, new sequencing datatypes (e.g. Illumina short-reads, PacBio long-reads or 10X barcoded reads) typically require new implementations of corresponding computational analysis techniques. Doing so requires software that is computationally efficient, quick to develop and easy to maintain in order to enable rapid adaptation to new kinds of data.

However, writing high-performance software for computational genomics, and for bioinformatics in general, remains a difficult task. Crafting an efficient software tool requires highly specialized domain expertise in performance engineering, computational modeling, and the ability to translate biological assumptions into algorithm and software optimizations [39]. As a result, most high-performance genomics software is highly application-specific and opaque, lacks appropriate documentation, and is difficult to maintain without a dedicated team of expert programmers. These factors collectively have sparked a replication crisis that is currently plaguing the field of computational biology and life sciences in general [30, 5, 20]. A lack of maintainability means that updating many software packages to work on modern types of data is difficult. For example, in many instances, an algorithm is optimized for performance on an old datatype (e.g. Illumina short-reads) and therefore cannot efficiently process a new datatype (e.g. IonTorrent short-reads, or PacBio or Oxford Nanopore long-reads). In general, there is a lack of good tools that can help researchers update old software or even develop new software; one typically has to choose between a software ecosystem that allows for rapid development at the expense of performance and scalability (e.g. Python or R ecosystems), or low-level languages that have acceptable performance but are hard to develop and maintain.

Here we present Seq (Figure 1), a domain-specific, high-performance programming language for bioinformatics and computational genomics, that bridges the gap between the ease-of-use and clarity of high-level languages like Python or MATLAB, and the performance of low-level languages like C or C++. Seq has the syntax and semantics of the widely-used, high-level Python language, on top of which it adds genomics-specific language constructs, types, and optimizations invisible to the user. Seq is to bioinformatics what MATLAB is to numerical computing—an accessible portal to the world of high-performance data science.

**Figure 1:**
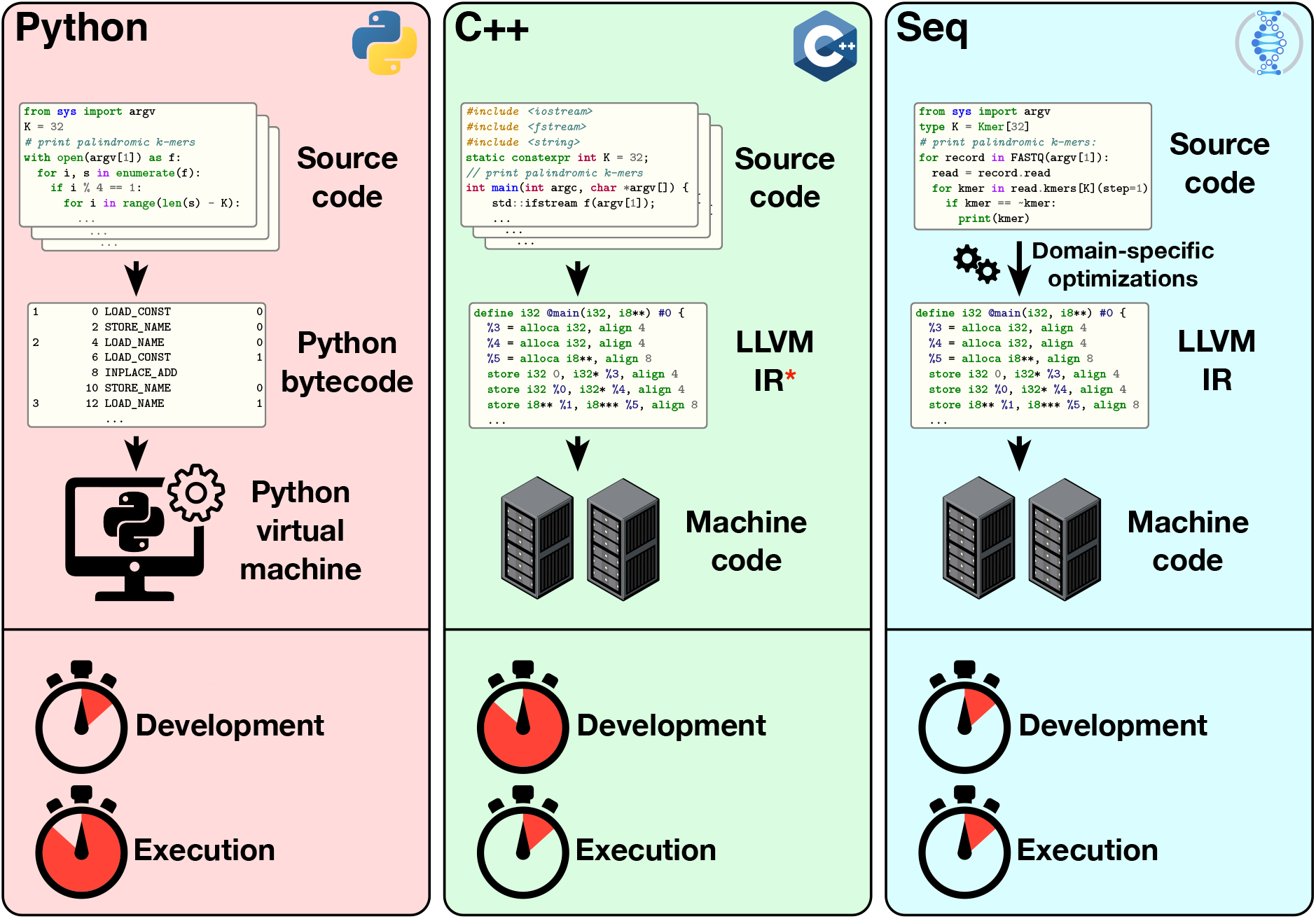
Overview of the Seq language and compiler, as compared to C++ and Python. The Seq language is an extension of Python with a multitude of genomics- and bioinformatics-specific language features, types and functionalities. The Seq compiler analyzes Seq source code and performs domain-specific optimizations that are hidden from the user, for sequence manipulation, memory management, genomic pipelines and more, whereupon the code is transformed to a LLVM (“Low-Level Virtual Machine”) Intermediate Representation, at which point the LLVM backend emits high-performance machine code. Seq marries the high-performance of low-level languages like C++ with the ease-of-use, clarity and maintainability of high-level languages like Python. *Note that not every C++ compiler emits LLVM IR; `clang` is among the most popular C++ compilers and uses an LLVM backend.

While some of these optimizations have been applied individually, no framework is currently able to seamlessly integrate them under a single, easy-to-use interface. To this end, we provide use cases where Seq improves on current approaches by re-implementing a myriad of popular genomics applications in the language, spanning tasks such as finding super-maximal exact matches (BWA-MEM [22]), genome homology table construction (CORA [38]), Hamming distance-based all-mapping (mrsFAST [16]), long-read alignment (minimap2 [23]), SAM/BAM preprocessing (GATK [26]), global sequence alignment (AVID [10]) and haplotype phasing (HapTree [9, 8]). We show that Seq’s compiler optimizations result in substantial performance improvements over these tools, including a 2× improvement in FM-index querying, up to a 4.5× improvement in Smith-Waterman alignment, up to a 3× improvement in Hamming-distance based read mapping, and up to an order of magnitude improvement in data preprocessing, genome index construction and numerical downstream analysis (**Table 1**)—all without loss of accuracy. Many of these optimizations are exceedingly difficult to implement by hand, and would require large-scale code changes and significant testing time to incorporate into existing applications; yet, in Seq, they often require just a single additional line of code. We also show how Seq can be used to rapidly re-implement an existing state-of-the-art tool from the literature (AVID) for which source code is not available—a useful capability in a time when many promising algorithmic ideas have no readily-available implementations. Taken together, we demonstrate how Seq can effectively optimize each stage of a typical genomics pipeline, from simple tasks like data parsing to more complicated tasks like dynamic programming or numerical optimization, while allowing for a clear, concise and maintainable codebase.

**Table .1:**
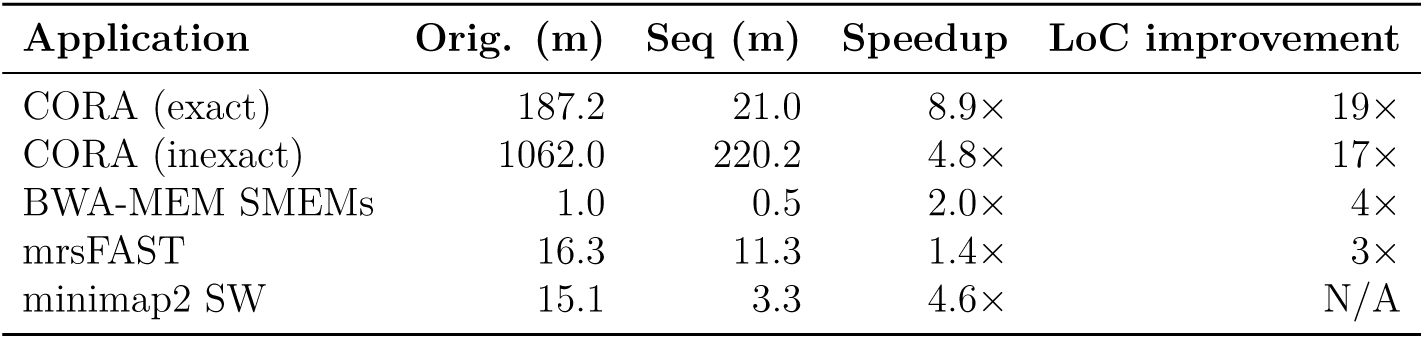
Overview of Seq’s performance (given in minutes) and “lines of code” (LoC) improvements on several applications.

**Table .2:**
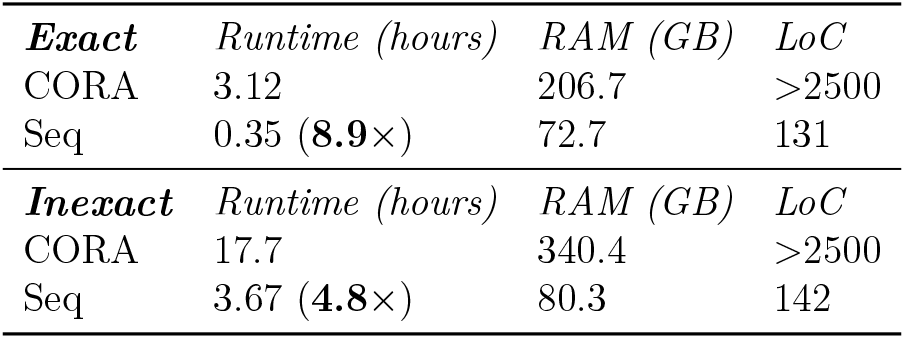
Comparison of original CORA homology table implementation (C++) and Seq implementation, in terms of runtime, memory usage and lines of code. Note that the original CORA implementation does not separate the exact and inexact implementations into separate modules, and uses the indicated number of lines for both in total.

**Table .3:** Comparison of SMEM-finding performance between BWA-MEM (timed using BWA’s fastmap command) and a Seq implementation with and without the prefetch optimization (code difference between Seq versions is one line – a single @prefetch annotation). Note that Seq’s FMD-index implementation does not use a sampled suffix array, leading to a larger memory footprint (similar to BWA-MEM2, which abandons the sampled suffix array for performance reasons).

|  | *Runtime (s)* | *RAM (GB)* | *LoC* |
| --- | --- | --- | --- |
| BWA-MEM | 62.2 | 4.6 | >500 |
| Seq | 50.5 (**1.2×**) | 48.7 | 133 |
| Seq + p/f | 28.8 (**2.2×**) | 49.0 | 134 |

**Table .4:** Comparison of original mrsFAST-Ultra with Seq implementation, for Hamming distance threshold *e* = 2 on chromosome 1 (top) and exact matching on the entire genome using an FM-index instead of a hash table as in the original (bottom). mrsFAST-Ultra was given 10GB for the read index in the chr1 experiment (`-mem` parameter); in the hg19 experiment, the first mrsFAST row shows a run for 15GB while the second shows a run with no memory limit—these values were chosen so as to make the memory footprints of both implementations roughly equal for a fairer comparison. The bottom row shows a Seq implementation that also uses the prefetch pipeline optimization (one line code difference).

|  |  |  |  |
| --- | --- | --- | --- |
| **chr1**, $e = 2$ | *Runtime (s)* | *RAM (GB)* | *LoC* |
| mrsFAST | 978.8 | 17.3 | >1000 |
| Seq | 677.3 (**1.4**×) | 15.5 | 373 |
| **hg19**, $e = 0$ | *Runtime (s)* | *RAM (GB)* | *LoC* |
| mrsFAST | 4,044.0 | 20.8 | >1000 |
| mrsFAST | 2,664.5 | 38.3 | >1000 |
| Seq | 1,310.2 (**3.1**×) | 22.2 | 99 |
| Seq + p/f | 922.3 (**4.4**×) | 22.2 | 100 |

**Table .5:**
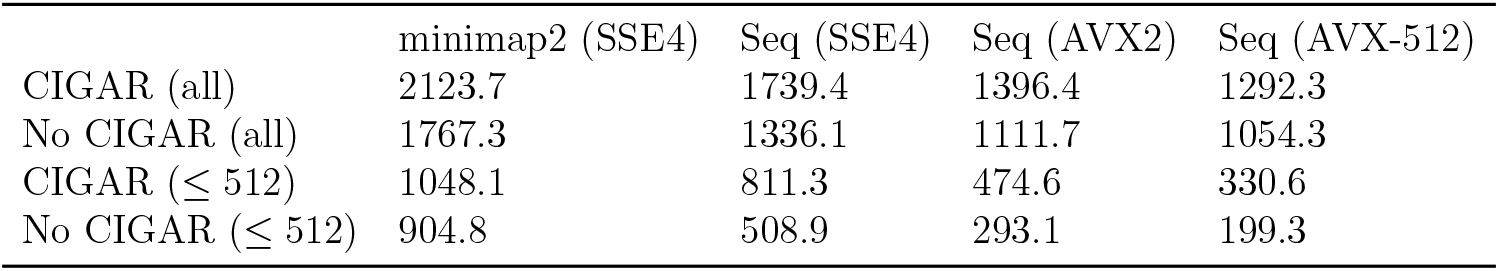
Runtime (in seconds) comparison of minimap2’s hand-vectorized intra-sequence alignment kernel (KSW2) with Seq’s inter-sequence alignment optimization, on various SIMD instruction sets. Performance was measured by aligning pairs of sequences that minimap2 aligns when run on a dataset of Nanopore long-reads.

|  | minimap2 (SSE4) | Seq (SSE4) | Seq (AVX2) | Seq (AVX-512) |
| --- | --- | --- | --- | --- |
| CIGAR (all) | 2123.7 | 1739.4 | 1396.4 | 1292.3 |
| No CIGAR (all) | 1767.3 | 1336.1 | 1111.7 | 1054.3 |
| CIGAR ($\leq 512$) | 1048.1 | 811.3 | 474.6 | 330.6 |
| No CIGAR ($\leq 512$) | 904.8 | 508.9 | 293.1 | 199.3 |

Conceptual advances include introducing new language features into Seq that allow for the seamless expression of complex genomics operations, algorithms and pipelines. We additionally introduce, implement, and evaluate domain-specific compiler optimizations related to fundamental genomics operations like reverse complementation, *k*-mer processing, Hamming distance computation, sequence alignment and large genomic index querying (e.g. FM-index querying). All of these optimizations are readily available out-of-the-box with Seq. We show how careful memory management, hidden from the user, allows for the efficient implementation of lengthy preprocessing steps intrinsic to every genomics pipeline. Seq enables everyday researchers—including those without a deep knowledge of computer science or programming—to rapidly write efficient software without the hassle of low-level implementation details or optimizations, which Seq handles automatically.

## Results

A typical genomics analysis pipeline currently consists of four fundamental types of stages: (i) data preprocessing, (ii) reference sequence processing, (iii) read sequence processing, and (iv) downstream analysis. Each of these stages employs different classes of algorithms and has distinct performance challenges (e.g. processing of reads often involves memory-bound operations whereas downstream analyses are frequently numerically-intensive). Thus, unique optimization techniques must be used at each stage in order to achieve ideal scalability and performance.

To demonstrate the utility of Seq, we re-implemented seven standard, real-world applications to evaluate the performance of the compiler’s domain-specific optimizations and the resulting code complexity across these stages, as compared to existing highly-optimized and widely-used C, C++, Java or Python implementations. Here we describe each of these applications in greater detail, as well as what is gained by porting them to Seq. A visualization of the genomics pipeline showing each of these applications is given in Figure 2.

**Figure 2:**
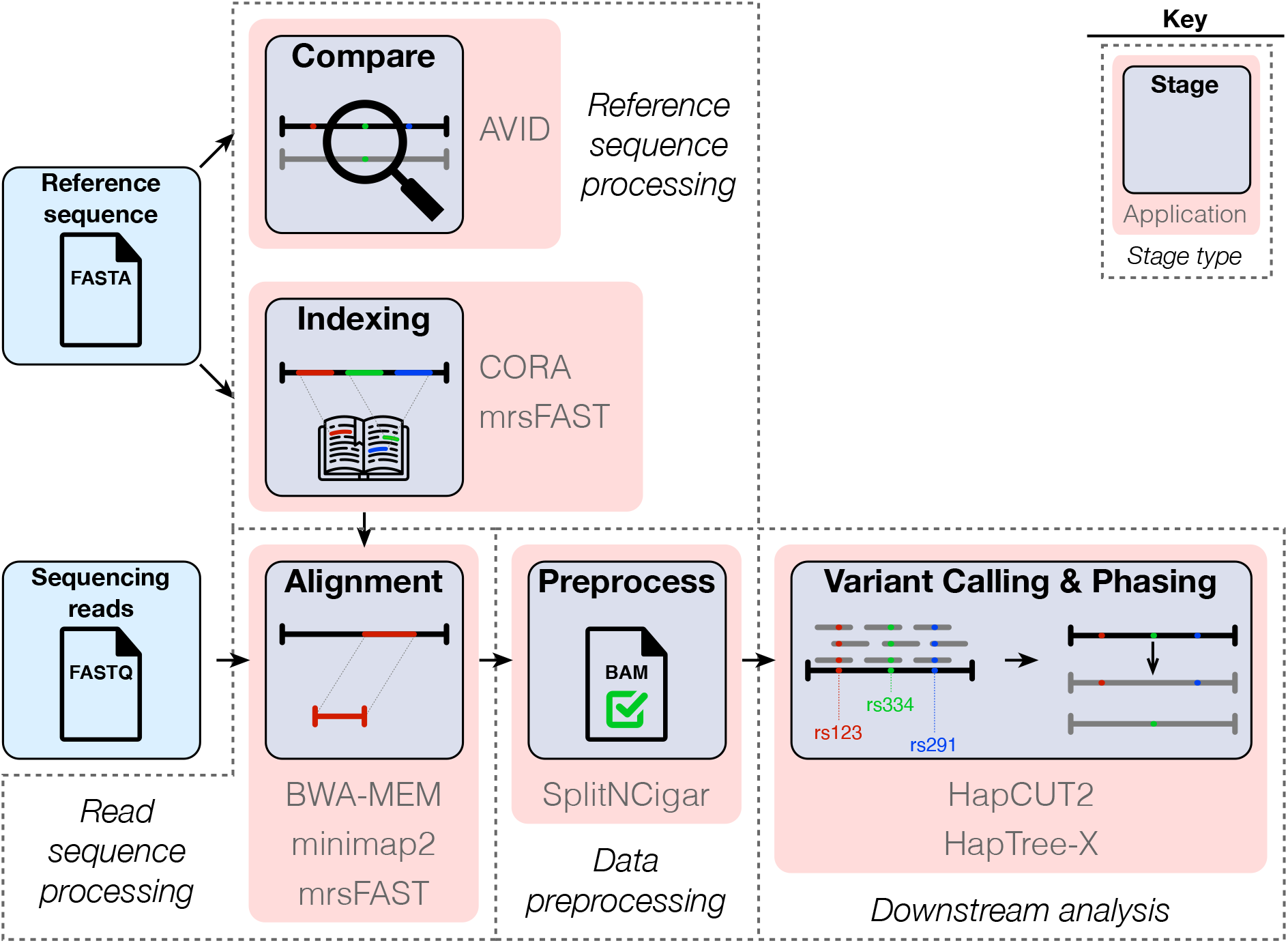
The standard genomics analysis pipeline, annotated with the state-of-the-art applications that we compare against here. The first step of most analyses involves indexing the reference to facilitate read alignment, which is done by CORA and mrsFAST (*reference sequence processing*). The alignment stage then maps reads to the reference, as is done e.g. by BWA-MEM, minimap2 and mrsFAST (*read sequence processing*). The resulting alignments (typically output as a BAM file) are preprocessed by tools like GATK’s `SplitNCigar` (*data preprocessing*), after which variant calling and phasing are performed by tools like HapCUT2 and HapTree-X (*downstream analysis*). Alternatively, analyses pertaining to comparative genomics involve comparing one or more reference sequences, which can be accomplished by global sequence alignment tools like AVID (also *reference sequence processing*).

### (i) Reference sequence processing

#### Genome homology table construction (CORA)

CORA [38] is an all-mapping tool for nextgeneration sequencing (NGS) reads. In other words, given a read (or a read pair), CORA reports all alignments (up to some edit or Hamming distance) of that read in a reference sequence. To achieve this goal, the primary data structure used by CORA is the *homology table*, which stores groups of “homologous” regions in the reference, and is used to convert a single alignment as reported by an off-the-shelf mapper like BWA, into a list of all mappings satisfying the distance criteria. Two versions of the homology table are built: *exact* and *inexact*. The exact homology table stores in a compressed form pairs of equal *k*-mers (with 33 ≤ *k* ≤ 64) from the reference, along with their loci (where “compression” is done by merging two or more consecutive homologous pairs into a single group). Similarly, the inexact homology table stores pairs of unequal *k*-mers whose Hamming distance is less than some threshold. CORA’s definition of homology also allows for cases where one *k*-mer is homologous to the reverse complement of another *k*-mer.

The process of constructing the homology table involves many of the operations that Seq optimizes, including *k*-merization, reverse complementation, *k*-mer matching and Hamming distance calculations. Indeed, we implemented CORA’s homology table construction in Seq, and compared both performance and lines of code with the original, highly-optimized C++ implementation. We used a *k*-mer length of 64 and a maximum Hamming distance threshold of 1 for the inexact homology table. Exact homology table construction was single-threaded, whereas the inexact table’s construction was allowed to use all 48 threads of our machine.

Figure 3a shows the results of the comparison. The Seq versions were substantially smaller in terms of lines of code (over 2,500 in the original C++ version versus less than 300 in Seq—nearly 10 ×fewer lines) and ran substantially faster (5–9×) while using 3–4× less memory. Lastly, we found the outputs of the original and Seq implementations to be very slightly different; a manual inspection revealed no issues with Seq’s output, but some incorrectly reported or missed homologies by the original implementation; therefore, we attribute this to a bug in the original version. Note that while such bugs are hard to find in large and tightly optimized C/C++ codebases, they often become self-evident in high-level codebases that focus only on the core algorithm, such as those produced with Seq.

**Figure 3:**
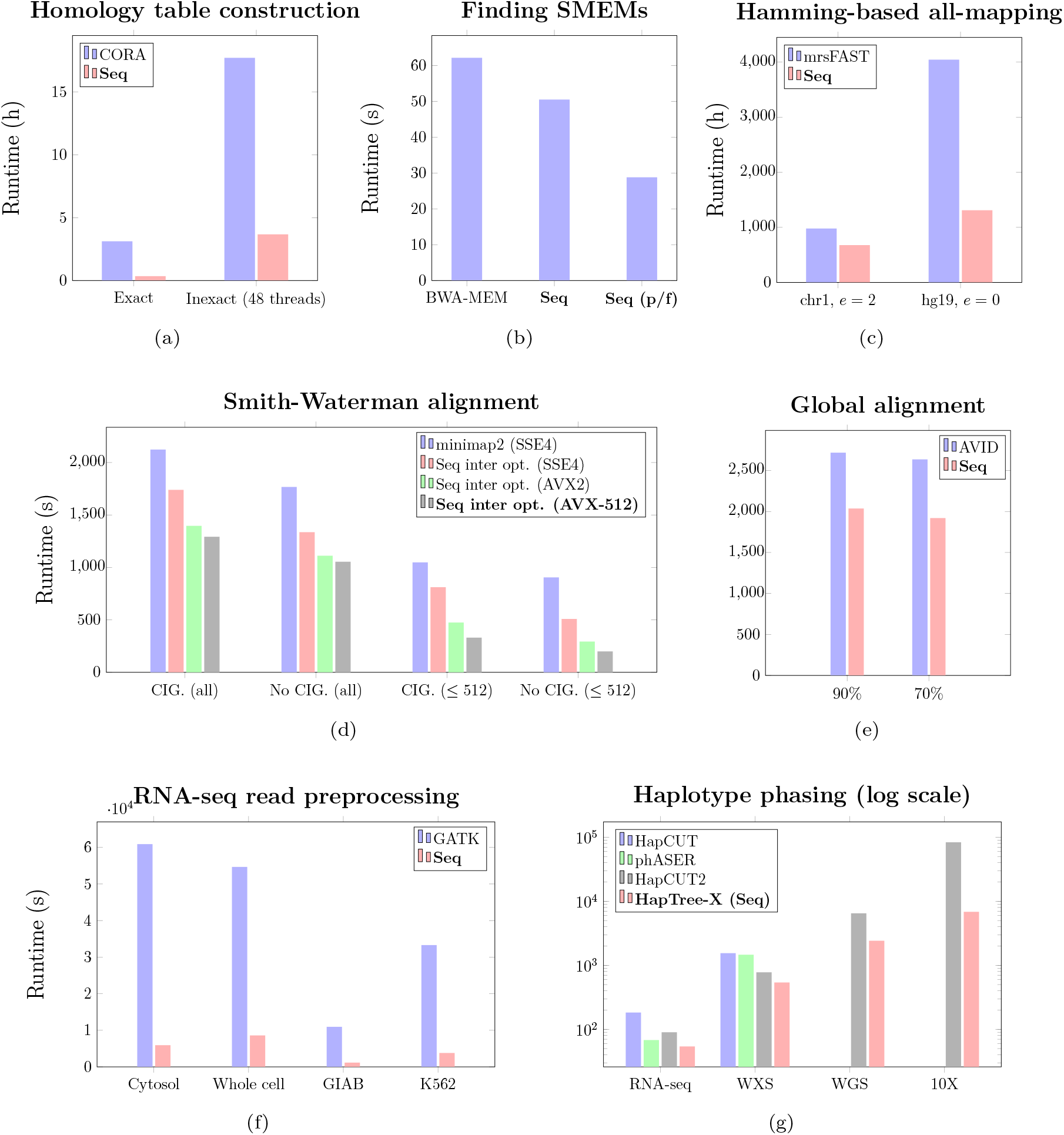
Performance improvements in seven standard applications from using the Seq language and compiler. The tested applications include genome homology table construction (from the CORA framework), finding super-maximal exact matches (i.e. SMEMs; from BWA-MEM), Hamming distance based all-mapping (from the mrsFAST mapper), Smith-Waterman alignment (from minimap2), global alignment (AVID), RNA-seq preprocessing (from GATK), and a novel algorithm for haplotype phasing compared to several state-of-the-art phasing tools. In the SMEMs benchmark, “Seq (p/f)” refers to Seq’s prefetch optimization. In the Smith-Waterman benchmark, “CIG.” refers to CIGAR string generation (i.e. the backtrace step), and the “≤ 512” experiments were run on sequences shorter than that limit. Seq’s inter-sequence optimization (“inter opt.”) was applied in the Smith-Waterman benchmark on several instruction sets (SSE4, AVX2 and AVX-512).

#### Global sequence alignment (AVID)

AVID [10] is one of the fastest tools for global alignment of large nucleotide and amino acid sequences (see also Batzoglou et al. [6]). Although the canonical global alignment algorithm (Needleman-Wunsch) is a slight variant of the local alignment algorithm (Smith-Waterman), only a few tools such as FASTA [29] and AVID (released in 1989 and 2002, respectively) are able to quickly produce global alignments between two sequences. Unfortunately, the source code of AVID is not available, and prebuilt binaries are only available for a limited number of platforms.

We re-implemented AVID’s algorithm as described in the original publication [10]. The core of AVID’s method consists of finding “Maximal Exact Matches” (MEMs)—identical sub-sequences between the two sequences of interest that cannot be further extended— and using them to guiding the global alignment process. MEMs are commonly found by recursively traversing a generalized suffix tree made from the subject sequences. In our implementation, we used suffix arrays (SA) instead of suffix trees (as Seq supports efficient SA operations out of the box), and a bottom-up traversal algorithm over SAs to find all MEMs [1].

Overall, we were able to prototype the whole pipeline as described in the paper in less than 200 lines of high-level Seq code. We ran AVID and Seq-AVID on the large set of similar (similarity ≥ 90%) and less-similar sequences (≥75%) from the human genome segmental duplications database [4, 27], and found that the Seq implementation provided up to a 5×speedup over the original binary while maintaining a similar accuracy (note that the exact algorithm used by the currently available AVID binary might differ slightly from the published description). The results of this comparison are shown in Figure 3e.

### (ii) Read sequence processing

#### Finding SMEMs (BWA-MEM)

Super-Maximal Exact Matches, or SMEMs, are a subset of MEMs such that no two SMEMs overlap within the read sequence. SMEMs are an integral component of many important genomics algorithms, ranging from sequence alignment to assembly [21, 22, 37]. Because the SMEM detection algorithm primarily involves querying an FMD-index data structure (unlike the MEM detection algorithm, which relies on suffix tree traversal), it is inherently memory-bound, and can be greatly accelerated by several of Seq’s compiler optimizations.

To this end, we implemented BWA-MEM’s SMEM-finding algorithm in Seq, with roughly 135 lines of code (compared to the over 500 lines of C code comprising the original). The standard algorithm involves repeatedly querying an FMD-index to extend matches between a read and the reference, which causes expensive memory stalls—indeed, this is the main bottleneck in the algorithm. As querying large genomic indices is itself a prevalent operation in genomics algorithms, the Seq compiler automatically performs several optimizations to greatly speed up such queries. Rather than issuing memory requests serially and forcing the entire program to stall, Seq makes use of coroutines and software prefetching to *overlap* the memory-bound stall from one query with other useful work, greatly reducing the effective stall time. This process is shown in Figure 7 (left half).

**Figure 4:**
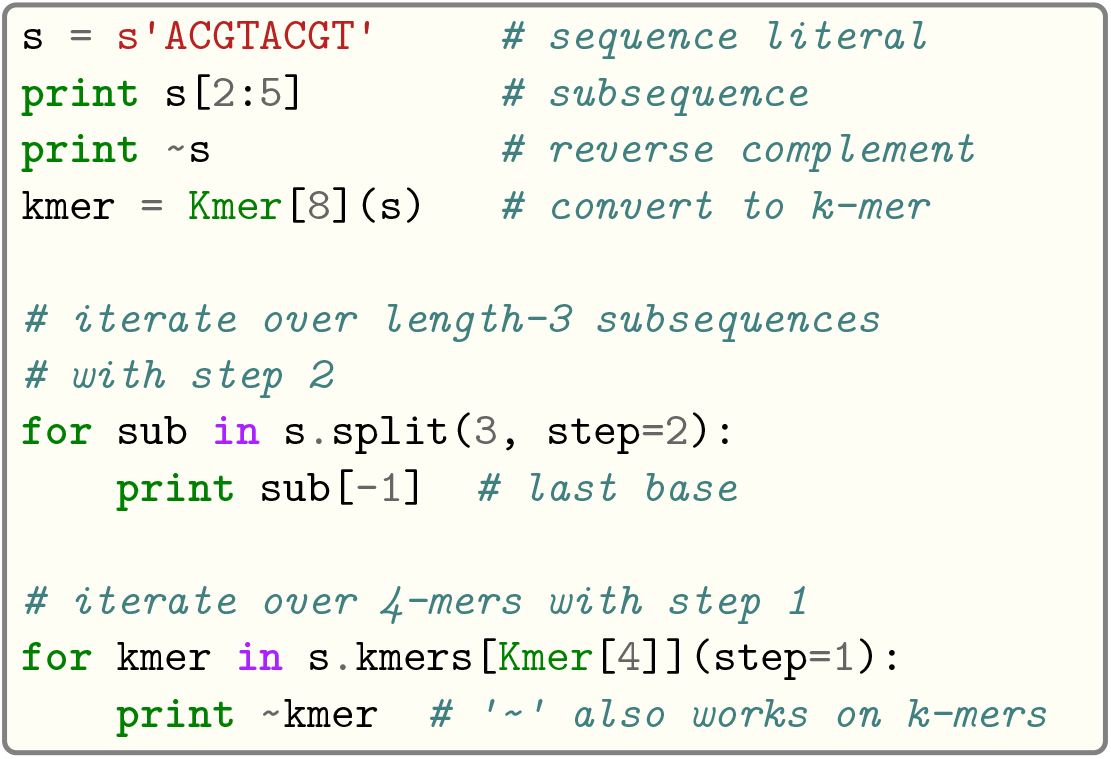
Examples of using sequence and k-mer types in Seq.

**Figure 5:**
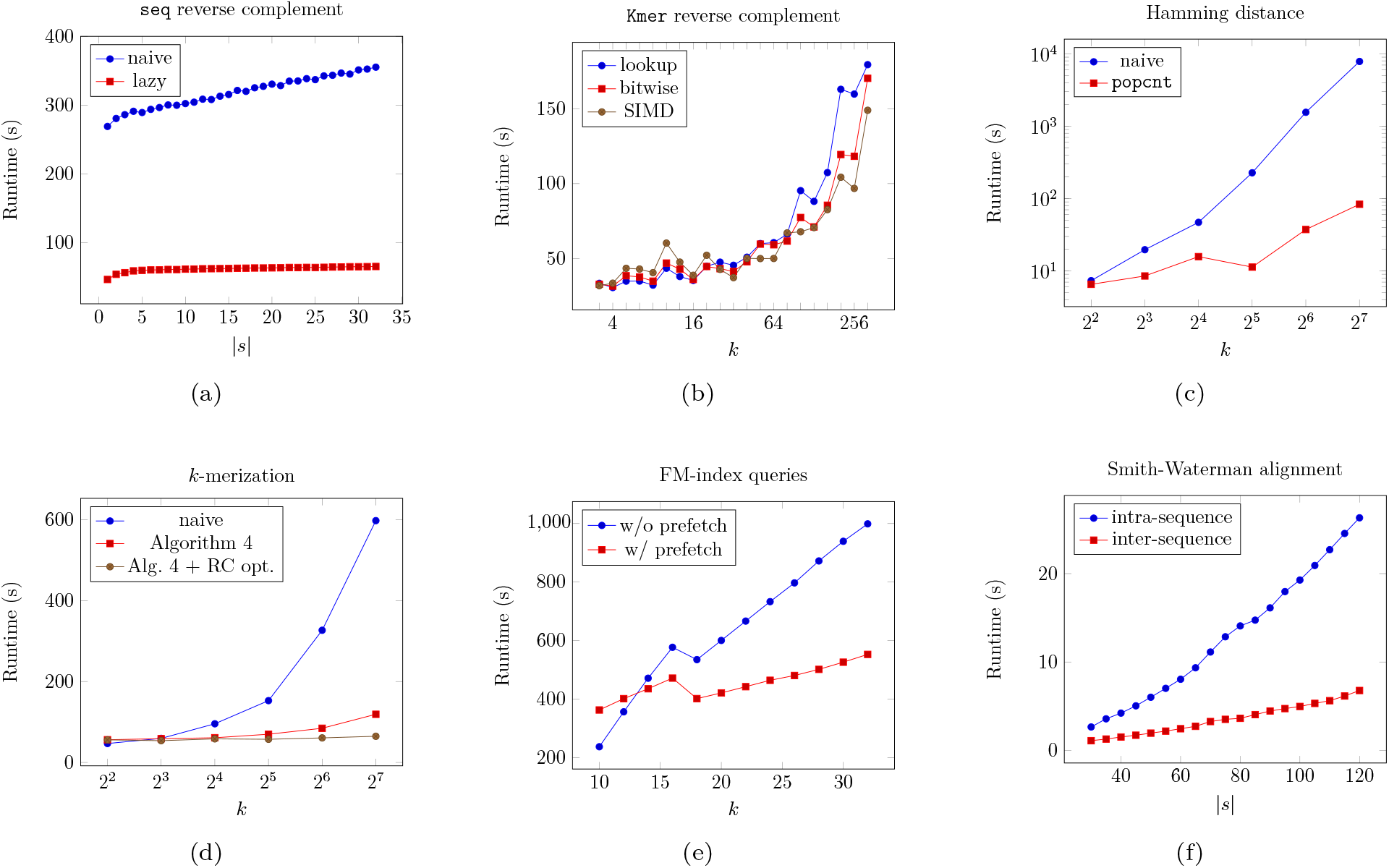
Effect of several compiler optimizations performed by the Seq compiler on various small benchmarks. Reverse complement performance was measured by counting sequences (or *k*-mers) lexicographically larger than their reverse complements; Hamming distance performance by computing distances between *k*-mers and their reverse complements; FM-index query performance by querying *k*-mers from a set of Illumina reads; Smith-Waterman alignment by aligning one million simulated sequence pairs of maximum length |*s*| with 90% identity and maximum indel size of 3. The reference genome used in each applicable benchmark was hg19.

**Figure 6:**
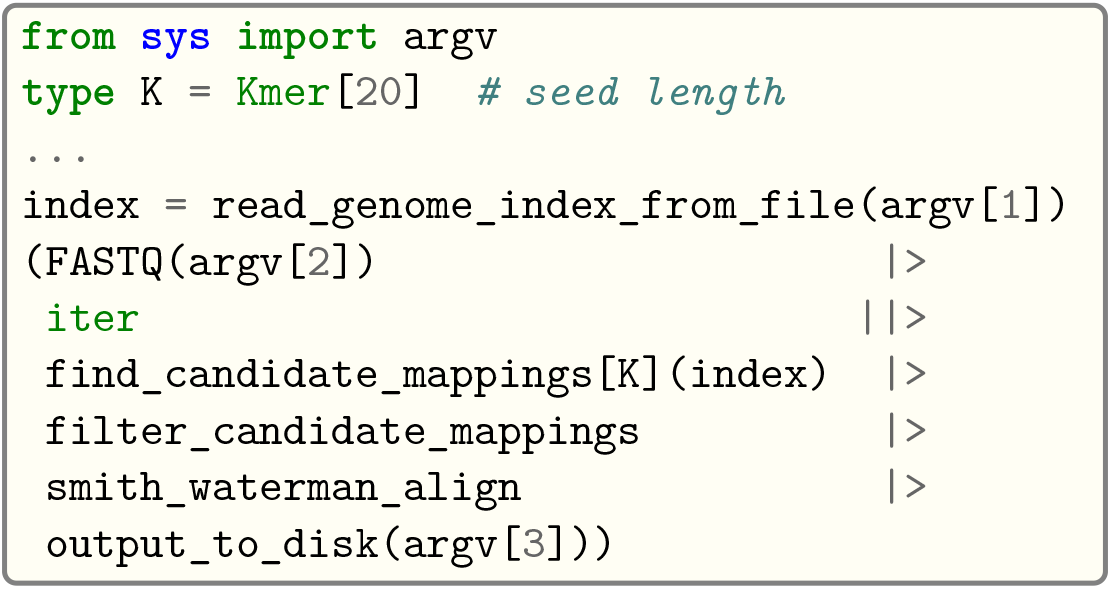
Hypothetical pipeline for read mapping in Seq, where the first argument (`argv[1]`) is some serialized genome index file, the second is a FASTQ file containing input reads, and the third is the output file. The shown pipeline is parallelized using the parallel pipe operator, `||>`, and each read can be processed in parallel.

**Figure 7:**
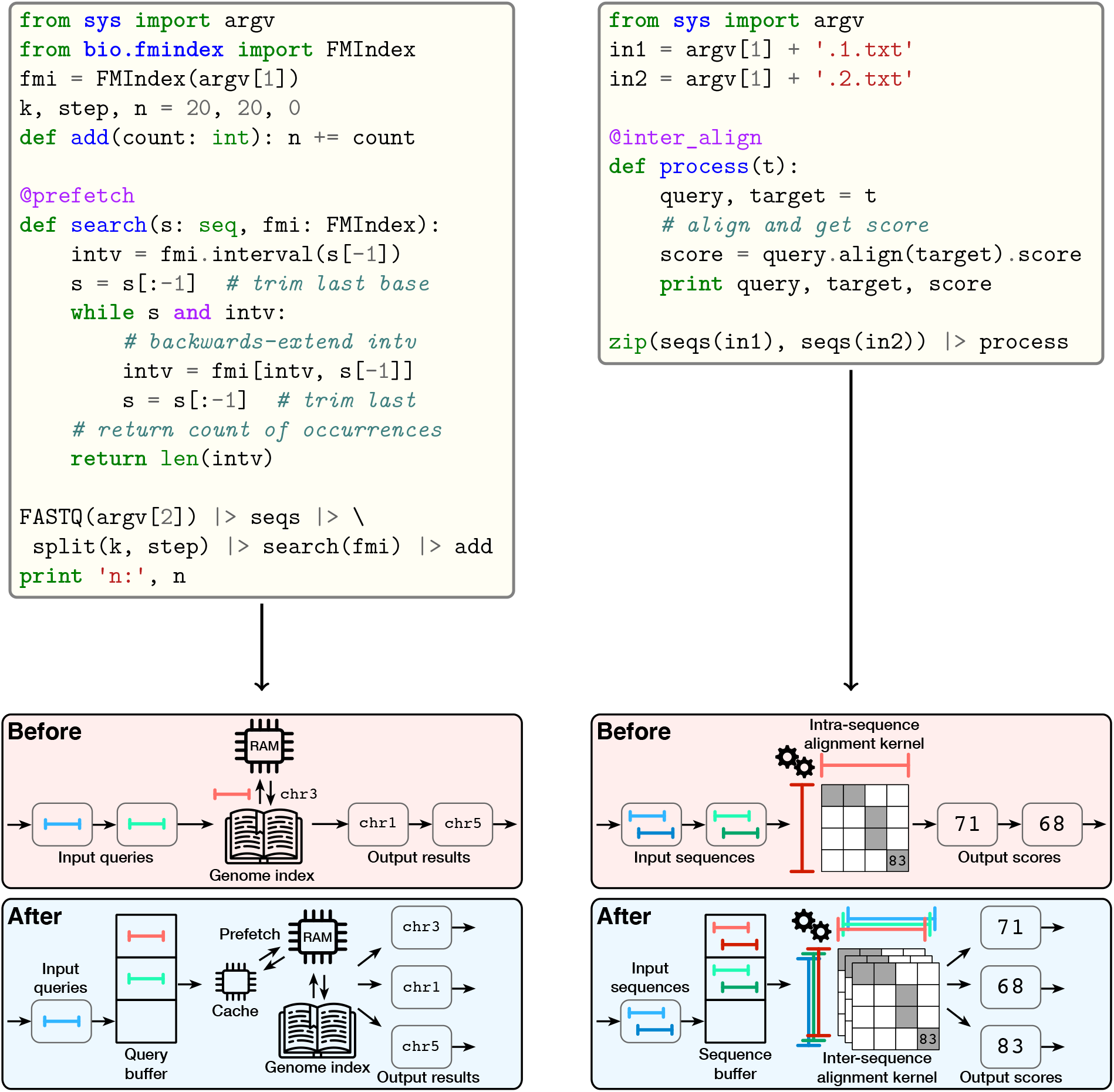
Coroutine-based pipeline optimizations in Seq. The top left snippet gives an example of using the @prefetch annotation with a pipeline performing FM-index queries; the depicted code simply counts occurrences of length-20 subsequences from a given FASTQ using the index. The top right snippet gives an example of inter-sequence alignment using the @inter_align annotation, where sequence pairs are read from two text files, aligned, and printed. The bottom row illustrates the resulting pipeline optimizations performed by the Seq compiler. For genomic index prefetching, pipeline stages that query a large index are converted to coroutines and suspended before querying the index, after issuing a corresponding software prefetch; a dynamic scheduler manages the coroutines and ultimately ensures that any cache miss latency is overlapped with useful work by another coroutine. For inter-sequence alignment, in the original pipeline, sequence pairs are passed through the pipeline one at a time and aligned individually; in the transformed pipeline, sequence pairs are batched by a dynamic scheduler, aligned with inter-sequence alignment, and returned to their respective suspended coroutines.

While implementing these transformations by hand in existing codebases would require substantial effort (including implementing coroutines, scheduling and managing coroutine state, dispatching queries and returning results, debugging, etc.), in Seq it only requires a single additional @prefetch annotation—everything else is handled automatically by the compiler. Applying this optimization to the SMEM algorithm resulted in a 2.2× speedup over BWA-MEM (Figure 3b).

Software prefetching is also used in BWA-MEM2, the next iteration of BWA-MEM, which adds numerous manual, low-level performance optimizations [37]. However, BWA-MEM2’s implementation does not perform a coroutine-based software context switch like Seq compiler does; consequently, as the authors describe, it cannot fully remedy memory-bound stalls. By contrast, Seq’s use of coroutines to overlap stalls with useful work can often ensure that a particular address is cache-resident by the time it needs to be accessed (i.e. when a suspended coroutine is resumed, as described in detail in Methods). We also evaluated software prefetching *without* the use of coroutines—much like what BWA-MEM2 does—but observed no appreciable performance improvements over the coroutine-based prefetching.

#### Hamming-distance based all-mapping (mrsFAST)

mrsFAST [16] is a perfect-sensitivity all-mapping tool based on Hamming distance. mrsFAST employs a hash table to index *k*-mers from a reference genome, and uses that table to find all mappings of a given read that are below a user-defined Hamming distance threshold *e*. The algorithm is centered around the pigeonhole principle: each read is partitioned into *e* + 1 non-overlapping seeds, which guarantees that at least one seed will map to all positions in the reference at which the Hamming distance is at most *e*. mrsFAST-Ultra, the latest version of the tool, is implemented in C, and involves many of the operations discussed here, including *k*-merization, reverse complementation, *k*-mer hashing, indexing and of course Hamming distance calculations.

We implemented a subset of mrsFAST-Ultra for single-end mapping in Seq to further test the effectiveness of the compiler’s optimizations, the results of which are shown in Figure 3c. With a distance threshold *e* = 2 on chromosome 1, the Seq implementation was 40% faster than the original hand-optimized C implementation, with a third of the code size. We also implemented a version using an FM-index instead of a hash table—a modification that requires changing only a few lines of code in Seq—and tested it with exact matching (*e* = 0) on the entire genome, resulting in a 3.1× improvement, or 4.4× with the prefetch optimization, which is particularly notable given that mrsFAST-Ultra’s is a cache-oblivious algorithm. Ultimately, the original version is comprised of well over 1000 lines of code and takes over an hour to perform exact matching, whereas the Seq version is comprised of 100 lines of code and completes in about 15 minutes.

#### Long-read mapping (minimap2)

As third-generation sequencing becomes more widespread, high-performance methods and tools for processing long-reads become increasingly important [34]. minimap2 [23] is the current state of the art in terms of long-read alignment, both in terms of performance and accuracy. The algorithm used by minimap2 follows similar steps as AVID, and involves chaining to find overlaps between reads and the genome, followed by dynamic programming alignment between the seeds of a chain to obtain the final alignments.

Smith-Waterman sequence alignment is an essential kernel in many genomics applications, and is consequently a heavily researched area [15, 32, 23]. Most hand-optimized implementations use instruction-level SIMD parallelism to accelerate the alignment of a single pair of sequences—an approach referred to as intra-sequence alignment. However, yet another approach is to use SIMD to accelerate aligning *multiple* pairs of sequences, referred to as inter-sequence alignment [32]. While inter-sequence alignment can be substantially faster than intra-sequence alignment, it is typically cumbersome to implement with general-purpose programming languages due to the need to batch sequences, manage state, dispatch alignment results and so on. In Seq, however, the compiler performs several pipeline transformations to convert standard alignment to inter-sequence alignment with just a single-line code annotation.

We implemented minimap2’s Smith-Waterman alignment step in Seq to test the compiler’s inter-sequence alignment optimization relative to minimap2’s SIMD-optimized intrasequence alignment kernel (in fact, Seq’s default alignment kernel is the same one used by minimap2; minimap2 does not support inter-sequence alignment, however). Results of this comparison are shown in Figure 3d, where Seq’s inter-sequence alignment optimization is up to 4.5× faster than the intra-sequence implementation. Note that the traceback step (required to produce CIGAR strings) is not vectorized in minimap2 nor Seq, meaning timings including CIGAR generation are expected to be comparatively strictly slower than those without CIGAR generation. Additionally, Seq automatically demotes sequences that do not benefit from inter-sequence optimization^2^ to intra-sequence alignment (i.e., the same method is used to align such pairs in both the Seq implementation and in minimap2), so consequently timings including “all” sequences will also be comparatively slower. Nevertheless, even when including CIGAR generation and all sequence pairs, Seq performs up to 60% better than intra-sequence alignment.

One distinct advantage of the Seq language over general-purpose languages is the ability to incorporate complex pipeline optimizations—like those for inter-sequence alignment and prefetching—with minimal code changes. For example, adding inter-sequence alignment to the current C implementation of minimap2 would require large-scale code refactoring to batch sequences (and other state information) before performing the alignment, and the complete rewrite of a complex SIMD-based alignment kernel, whereas in Seq all of this is handled implicitly by the compiler using coroutines, and usually requires only a single-line code annotation, as shown in Figure 7 (right half).

### (iii) Data preprocessing

#### RNA-seq data clean-up (GATK SplitNCigar)

The most time-consuming steps in a NGS processing pipeline often consist of mundane data transformations performed on large datasets prior to, during and after the alignment or downstream analysis steps [28]. These tasks typically employ string operations (e.g. duplicate marking, sample tagging), score recalculations (e.g. base quality score re-calibration), and in the case of RNA-seq samples, splitting reads that span large intronic regions. Such tasks are commonly performed by GATK and Picard tools [11, 26]. However, these tools leave a lot to be desired in terms of performance, mainly due to the high-level language they are written in (Java). While data preprocessing tasks are often conceptually simple, the sheer volume of data (often on the order of terabytes) heavily penalizes even minor performance shortcomings that high-level languages often bear (e.g. an unnecessary memory copy can easily add several additional hours to the total runtime on real-world datasets). For instance, a relatively simple tool for splitting intron-spanning reads—`SplitNCigar`—from the GATK suite takes more than 15 hours to complete on a medium-sized 40GB BAM file.

We show that such preprocessing can be done faster and easier by re-implementing GATK’s `SplitNCigar` tool in Seq. By relying on automatic, conservative memory management and thus avoiding unnecessary memory copy operations, we were able to obtain 10× faster runtimes while using 10–20× less memory than the original GATK implementation (note that both Java and Seq rely on a garbage collector for managing memory). In concrete terms, this means that a 17-hour preprocessing step can be completed in just 100 minutes using Seq. Results of this comparison are shown in Figure 3f.

### (iv) Downstream analysis

#### Haplotype phasing (HapTree-X)

Haplotype phasing is the process of separating variants into a set of maternal and paternal alleles, and can provide valuable biological insights beyond what can be gained from raw genotypes alone [2]. HapTree-X is a phasing algorithm implemented in Seq on top of HapTree [9], which utilizes Bayesian optimization to infer optimal haplotypes from aligned genomic data. In contrast to other case studies that focus mostly on discrete optimization and sequence operations, the HapTree-X phasing algorithm is a probabilistic optimization framework that spends most of its runtime doing numerical computations and graph traversals. Furthermore, HapTree-X is a complete genomics analysis pipeline that needs to preprocess and analyze large BAM files prior to the haplotype phasing step. As such, it is an excellent test of Seq’s versatility and applicability to applications that require both efficient sequence processing *and* high-performance numerical algorithms.

Performance of Seq/HapTree-X on four diverse datasets spanning multiple sequencing technologies, relative to another three state-of-the-art phasing tools, is shown in Figure 3g. HapTree-X was up to 3× faster than phASER (Python) and HapCUT (C), and 1.4× faster than HapCUT2 (C++) on smaller RNA-seq and medium-sized exome datasets. However, we found that the performance gains increased with the scale of the data: on a large 120GB WGS BAM, HapTree-X was 3× faster than HapCUT2, while on a high-coverage 10X Genomics dataset (≈200GB BAM file), HapTree-X achieved up to a 10× speed up over Hap-CUT2 (C++, Python). Furthermore, parallelizing the HapTree-X pipeline (by simply converting a few nested function calls into a Seq pipeline) further decreased the total runtime by 2×. In absolute terms, HapTree-X was able to phase a 200GB file in less than an hour as compared to the 22 hours needed by the fastest alternative tool.

## Discussion

The creation of high-performance, maintainable bioinformatics software is undoubtedly a difficult task, as evidenced by the field’s unfortunate track record of poor software development, maintenance and testing practices. A key cause of this, among other factors, is a lack of domain-specific development tooling, debugging facilities and general programming infrastructure. While several software libraries have been developed as an attempt at answering this problem, libraries are unable to perform many of the optimizations discussed in this work and hence cannot achieve optimal performance in many key situations. Additionally, these libraries are still bound to their host languages, usually at the detriment of either performance or software maintainability.

The fundamental difference with Seq is that its entire software stack, from the language itself to the standard library to the compiler and even runtime, is designed and implemented with the domain of bioinformatics in mind. “Owning” the entire stack in this way comes with numerous distinct advantages. For example, datatypes or compiler optimizations relevant to new sequencing platforms can be incorporated seamlessly into the framework, circumventing large code rewrites or potentially even algorithmic changes. Additionally, as GPUs, FPGAs and cloud computing become increasingly prevalent in the field, new compiler backends for Seq—including new compiler optimizations pertinent to each computing platform—would enable existing code to run without changes on these backends, and is ongoing work within the Seq project. Indeed, many novel, promising computer architectures such as memory-driven computing [7] require specific optimizations and program transformations to be fully taken advantage of; these can be performed automatically by the Seq compiler. Yet another unique advantage of a framework like Seq lies in its ability to provide domain-specific debugging or visualization support; as the compiler has knowledge of sequences, *k*-mers and operations involving them, debugging or visualization tools can be seamlessly integrated into the Seq framework, allowing users to track and observe in a meaningful way how data is transferred and manipulated throughout a program. None of these possibilities are easily attainable without having a software stack designed from the ground up for genomics and bioinformatics. In fact, such domain-specific languages are widely used in a variety of other fields for similar reasons, including computer graphics [31], tensor algebra computing [18] and physical simulations [19].

We have presented the first step towards a rich ecosystem of inter-operable bioinformatics software development tooling, debugging support and development environments through a unified, expressive language and compiler that can act as the centerpiece of this ecosystem. By giving bioinformaticians, genomicists and researchers of all backgrounds a scalable way to prototype, experiment and analyze large biological datasets through a familiar and easy-to-use interface while leaving optimizations and performance considerations to the compiler, we hope that Seq will act as a catalyst for scientific discovery and innovation.

## Methods

The fundamental datatype underlying most bioinformatics or genomics applications is the “sequence”, which is conceptually a string of nucleotide (“A”, “C”, “G”, “T”) bases (or possibly amino acids in the case of protein sequences; while we focus on DNA sequences here, the same principles can be carried over to RNA or protein sequences, which Seq also has full support for). Here, we lay out how Seq represents and manipulates sequences and *k*-mers to optimize genomics applications. We also discuss the notion of a “pipeline”, including how Seq represents pipelines at the language level as well as the numerous pipeline optimizations the compiler performs.

### Preliminaries

For a sequence *s* over an alphabet ∑ (where usually ∑ = {`A, C, G, T`}), we let |*s*| be the length of *s, s*[*i*] be the character at 0-based index *i, s*[*i* : *j*] be the subsequence from 0-based index *i* (inclusive) to 0-based index *j* (exclusive), and *s* || *t* be the concatenation of *s* with another sequence *t*. Further, let 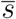 be the *reverse complement* of *s*. Reverse complementation is a common operation in computational genomics where a given sequence *s* is transformed into a new sequence *t* such that *t*[*i*] = RevComp(*s*[|*s*| − *i* − 1]) for 0 ≤ *i* < |*s*|, where

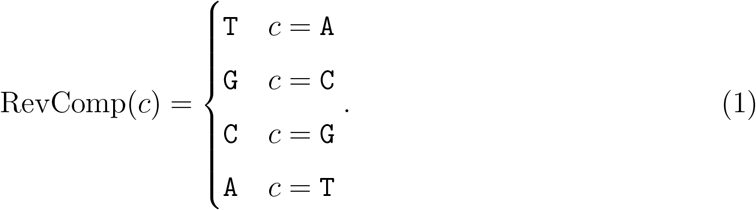

We define a *k*-mer to be a sequence *s* of fixed length |*s*| = *k*. *k* is typically on the order of 10 to 100. Let *κ_k_*(*s*) be the set of all *k*-mers that exist as subsequences in sequence *s*.

These definitions give rise to various algebraic rules, each of which Seq uses to perform various optimizations. For example:

1. 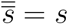: “the reverse complement of the reverse complement is the original sequence” – enables efficient *O*(1) lazy sequence reverse complementation.
2. 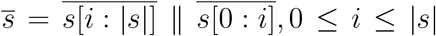: “the reverse complement is the concatenation of the reverse complements of the two halves of the sequence, in reverse order” – enables efficient *k*-mer reverse complementation via a lookup table.
3. 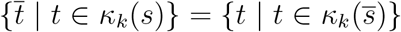: “iterating over reverse complemented *k*-mers can be done by iterating over *k*-mers of the reverse complemented sequence” – enables loop reordering for *k*-mer iteration followed by reverse complementation.

Below, we discuss in greater detail how each of these types and operations is used and implemented in Seq, and provide numerous benchmarks as to their efficacy.

### The Sequence Type

It goes without saying that computational genomics applications largely deal with processing and operating on sequences, and so an efficient *sequence type* implementation is invaluable. Seq provides a sequence type `seq` and a separate *k*-mer type `Kmer[`*k*] for 1 ≤ *k* ≤ 1024. While the sequence type and the *k*-mer type are conceptually both strings of nucleotide bases, they differ largely in where and how they are used: sequences have arbitrary length and can be accompanied by various metadata like an error profile, quality scores or ambiguous bases; *k*-mers on the other hand, at least in how they are frequently used in practice, are fixed-size segments of some larger sequence. For example, a typical alignment algorithm would employ short *k*-mers obtained from an arbitrary-length sequencing read, in conjunction with some *k*-mer index, to obtain candidate alignments for that read, where *k* is usually chosen as a fixed parameter beforehand, but read lengths are not known until runtime. In this case, the reads would be of type seq and the *k*-mers would, unsurprisingly, be of type `Kmer[`*k*] for some *k*. In practice, general sequences can also have a larger alphabet than *k*-mers (which are restricted to just the four nucleotide bases); for instance, `N` indicates an ambiguous base, `R` indicates an `A` or a `G`, etc. Figure 4 gives a few examples of using sequence and *k*-mer types in Seq, showing the general syntax as well as some common operations and how to convert between the two.

The following operations on sequences and *k*-mers are common, and are the subject of several compiler optimizations performed by Seq: *subsequence extraction/iteration, reverse complementation, k-merization* (the process of iterating over a sequence’s constituent *k*-mers), *Hamming distance computation, hashing / indexing*, and *dynamic programming alignment* (e.g. Smith-Waterman or Needleman-Wunsch).

### Implementation

Seq’s sequence type is implemented as a 16-byte pass-by-value structure containing an 8-byte character pointer and an 8-byte length. The C equivalent would be:

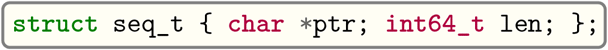

Subsequence operations do not copy, but rather just return a new sequence instance whose pointer is at the appropriate offset from the original’s. Seq then uses a conservative garbage collector to ensure that memory is deallocated as needed, and that nothing is prematurely deallocated, which is especially important when employing this scheme as many pointers will be referring to the middle of some larger allocated block. (In general, there are many ways to implement a sequence data type, and future work entails choosing the best variant based on context at compile-time, be it an ASCII representation, a 2-bit encoding or something else. We found the described implementation to work well in a wide range of cases, however.)

Sequence reverse complementation is an *O*(1) operation in Seq, implemented not by copying and physically reverse complementing, but by simply, “lazily”, flipping the sign of the sequence length. Then, each sequence method (e.g. for subsequence, *k*-merization, etc.) first checks the sign of the length, and uses it to determine if the sequence should be treated as reverse complemented or not (of course, the actual length is given by the absolute value). Conceptually, therefore, each sequence *s* is a 2-tuple of the underlying “logical” sequence *s*_log_ and a length *s*_len_, such that |*s*| = |*s*_log_| = |*s*_len_|, and *s*_len_ < 0 if and only if *s* is really reverse complemented with respect to *s*_log_. The reverse complementation algorithm is shown in Algorithm 1.

**Algorithm 1:**
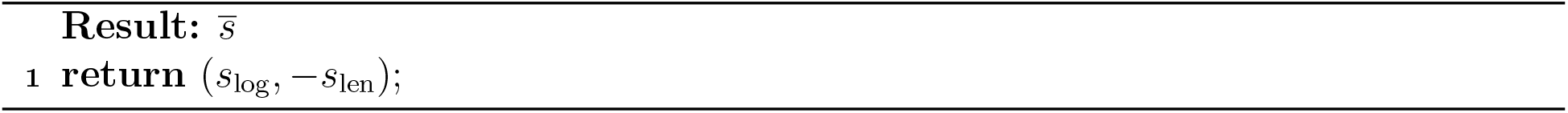
Reverse complement of sequence *s* = (*s*_log_, *s*_len_).

An example implementation of *s*[·] under this scheme is shown in Algorithm 2; notice that the sign of *s*_len_ must first be checked in order to determine how to index into *s*_log_. If *s*_len_ is positive, we index regularly; if it is negative, on the other hand, we treat *s*_log_ as reverse complemented and index correspondingly.

The appeal of this approach is that it not only saves memory and copying, but can also be applied multiple times without issue. For instance, clearly 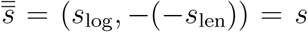. A more complicated example is 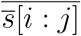, which also works as shown below, along with a concrete example where *s* = AAAGGGTTTCCC (|*s*| = 12) with *i* = 2 and *j* = 5:

**Algorithm 2:**
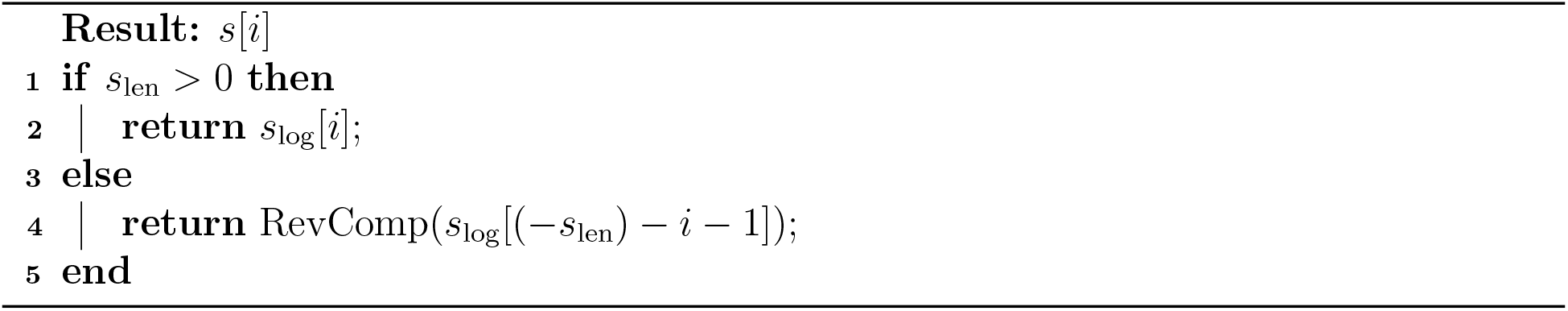
Sequence indexing for sequence *s* = (*s*_log_, *s*_len_).

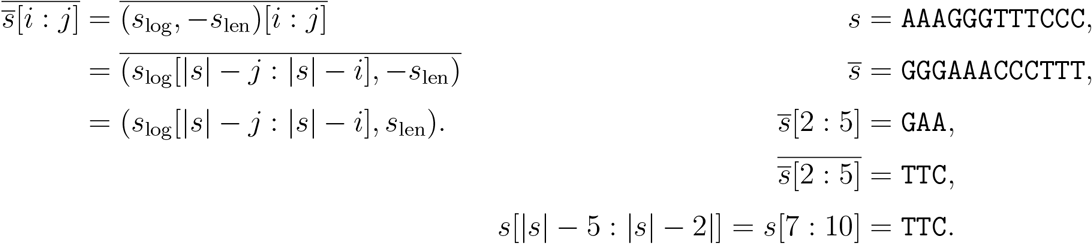

The drawback of this approach is that all sequence operations must include a preliminary check of *s*_len_’s sign, although we have found the overhead of this to be minimal, and greatly outweighed by the benefits (Figure 5a). In fact, this check can be elided altogether by the compiler in many cases (e.g. when sequences are first read from disk, at which point nothing has yet been reverse complemented).

Figure 5a shows the performance of this approach compared to the naive method of simply copying and reverse complementing. The benchmark shown in Figure 5a entails counting all subsequences of a given length in the human reference genome that are lexicographically larger than their reverse complement. Notice that the naive implementation is *O*(|*s*|) in the best case, whereas the lazy approach is best-case *O*(1) since bases must only be read until the lexicographic order of the given sequence and its reverse complement can be determined (by contrast, the naive approach must always read all bases of the subsequence to reverse complement it). Hence, the lazy approach not only performs better, but also scales better, as shown in the figure.

### k-mers

In Seq, a *k*-mer’s length is defined by its type (e.g. `Kmer[5]` is a different type than `Kmer[32]`), so unlike sequences the length does not need to be stored explicitly. Seq stores *k*-mers in a 2-bit encoded format, where `Kmer[`*k*] maps to LLVM IR type `i`*N* with *N* = 2*k*. LLVM then maps that integral type to hardware; for *k* ≤ 32, a *k*-mer fits in a single machine register on 64-bit systems, whereas larger *k*-mers may be spilled to the stack. Notationally, let *ξ_k_*(·) be the bijection from the set of all *k*-mers to unique 2-bit encodings.

### Reverse complement

The Seq compiler uses three different approaches to reverse complement *k*-mers:

- *Lookup*: The lookup method uses a lookup table of hard-coded 4-mer reverse complements (taking up 4^4^ × (2 · 4) bits, or 256 bytes). Reverse complements of longer *k*-mers are then constructed using the fact that 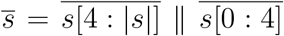 (recursively), i.e. the first four bases are reverse complemented, followed by the next four and so on, after which they are concatenated (in this case via bitwise shifts) in reverse order. For *k* < 4, the 2-bit encoding can simply be padded with `A` bases to construct a 4-mer, and the resulting `T` bases can be trimmed after reverse complementation.
- *Bitwise*: The bitwise approach uses a series of bitwise operations to reverse the 2-bit pairs of the encoded *k*-mer, then applies a bitwise-NOT to obtain the complement, the end result of which is the reverse complement. This is a generalization of the approaches used in GATB [14] and Jellyfish [25] for arbitrary-width integer encodings.
- *SIMD*: The SIMD approach uses a vector shuffle instruction to reverse the bytes of the encoded *k*-mer, then uses bitwise operations on each byte of the reversed vector in a similar fashion as the bitwise approach to obtain the final reverse complement. A similar method is implemented in MMseqs2 [35], although this implementation uses x86-specific pshufb instructions to reverse complement individual bytes in the reversed vector, whereas Seq’s LLVM IR implementation uses just the target-agnostic LLVM IR `shufflevector` instruction followed by a series of shifts and bitmasks to achieve the same effect.

The performance of each of these approaches for a wide range of *k* is shown in Figure 5b. As shown, there is no clear winner between these three methods. For smaller *k* (≤ 20), the lookup method is almost always best; for larger *k* (≥ 32), the SIMD approach is almost always best. For *k* values between these two cutoffs, the optimal algorithm varies, but we found the bitwise approach to be consistently close to, if not the best option. Because of this, the Seq compiler employs this heuristic to decide how to reverse complement a given *k*-mer.

### k-mer hashing

*k*-mer hashing is an extensively researched area [24]. For *k* ≤ 32, Seq simply uses the encoded 2-bit value as the hash by default. For *k* > 32, on the other hand, the situation is somewhat more complicated. One option is to use the hash of the first 32 bases, but due to the highly non-uniform structure of the genome, this leads to excessive collisions; for example, all *k*-mers consisting of a string of 32 `A`s followed by unique bases would hash to the same value. For this reason, hashing *k*-mers for *k* > 32 is done by hashing the first and last 32 bases, and XORing the two hash values. Compared to the naive approach, this method reduces the average collisions per unique 64-mer in hg19 from 1.089 to 1.007, and more than halves the size of the largest bucket. The best way to hash *k*-mers can vary based on the application, so Seq enables the user to override the *k*-mer hash function if needed.

### Hamming distance

The Hamming distance of two *k*-mers *s* and *t* is defined to be |{*i* | *s*[*i*] ≠ *t*[*i*], 0 ≤ *i* < *k*}|; i.e. the number of positions at which corresponding bases differ. A naive Hamming distance algorithm is to iterate over the bases of the two *k*-mers one at a time, and count how many differ. Seq employs a different algorithm using the `popcnt` instruction (which computes the number of 1 bits in an integer’s binary representation), shown in Algorithm 3.

**Algorithm 3:**
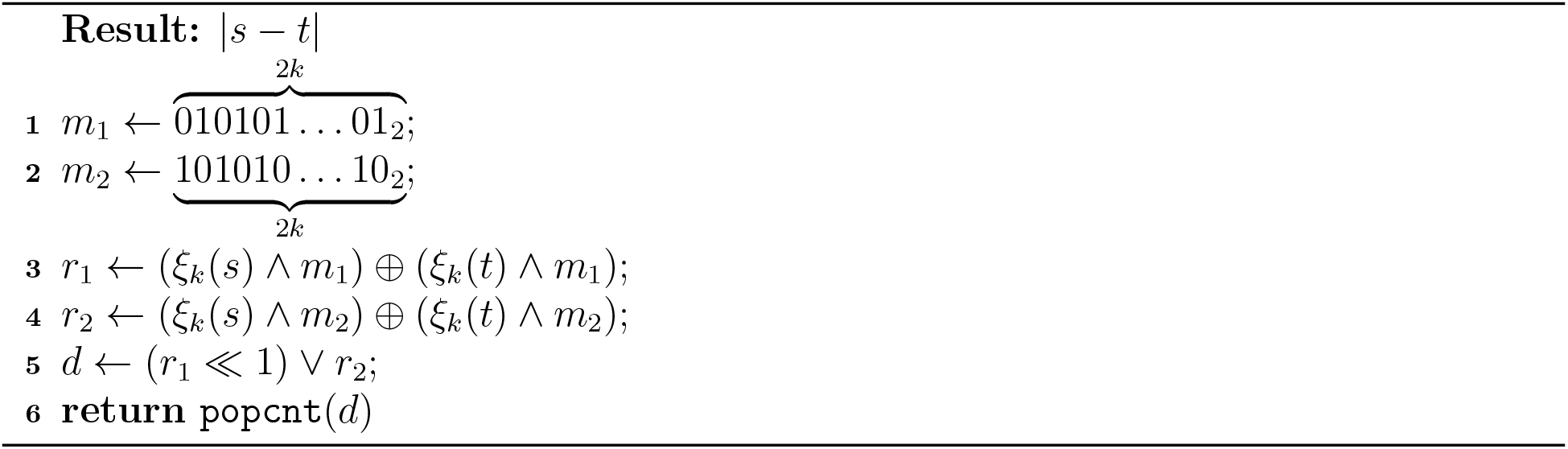
Hamming distance between two *k*-mers *s* and *t*.

Algorithm 3 works by first checking for differences in the odd bits via the *m*_1_ bitmask, then checks the even bits using *m*_2_, and finally ORs the results to obtain a bit-vector with 1s corresponding to different bases, a population count of which is the desired result. We leave to LLVM the decision of how to lower the logical `popcnt` instruction to actual machine instructions. For longer *k*-mers, the `popcnt` version can be substantially faster than the naive version, as shown in Figure 5c. For example, for 128-mers, the `popcnt` approach is nearly 100× faster.

### Pipelines

Pipelining is a natural model for thinking about processing genomic data. For example, a typical read mapping algorithm can be formulated as the pipeline:

1. Read input reads from FASTQ file
2. Extract seeds and lookup in some genomic index
3. Filter candidate alignment positions
4. Perform full Smith-Waterman alignment on remaining candidates
5. Format and output results to SAM file

Pipelining is supported natively in Seq via the pipe operator, `|>`. Pipelines can be parallelized via the parallel pipe operator, `||>`, which allows all subsequent stages to be executed in parallel (the Seq compiler uses an OpenMP task backend to implement this; future work entails implementing other backends, such as GPU for highly parallelizable tasks or MPI for distributed computing). In Seq, `a |> b` simply passes the output of `a` to `b` if `a` is a function; if `a` is a generator, all values yielded by `a` are passed to `b`. An example of how pipelines might be used to implement a typical read mapping application with the stages described above is shown in Figure 6.

The Seq compiler performs various domain-specific optimizations on pipelines, which we describe here.

### k-merization

An extremely common pattern is to iterate over the *k*-mers of some sequence with a particular step size *r*. In Seq, this can be achieved using the `kmers[K]` function, where `K` is the target *k*-mer type. For instance, the following code:

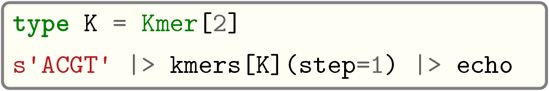

prints all 2-mers in the sequence `ACGT` (i.e. `AC, CG` and `GT`). Obtaining *k*-mers with positions as a tuple can be done with the analogous `kmers_with_pos[K]` function.

The best way to perform this *k*-merization depends on *k* and the step size *r*. For example, if *r ≥ k*, the best we can do is to simply extract the length-*k* subsequences at offsets 0, *r*, 2*r*, etc. and convert them to *k*-mers. However, if *r < k*, then *k*-mers overlap, so we can reuse the previous *k*-mer to obtain the next, similar to a rolling hash. The situation is complicated somewhat by the presence of non-`ACGT` bases in the sequence; *k*-mers spanning such bases are skipped by `kmers`. A simplified version of the *k*-merization algorithm used in Seq is shown in Algorithm 4 in the Appendix. In this algorithm, the two main branches inside the loop dictate whether the current *k*-mer must be “refreshed” because a non-`ACGT` base was encountered; if so, the *k*-mer is re-encoded afresh via SequenceToKmer, otherwise the previous *k*-mer is “shifted” to the right via ShiftKmer to cover the new bases, without doing a full re-encoding from scratch. Also notice that if *r ≥ k*, the second branch is never entered, and we perform a full re-encoding for every *k*-mer.

### Reverse complementation

*k*-merization followed by reverse complementation can be implemented more efficiently by using property 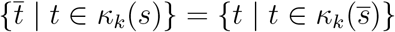, as described above. Because of this, the Seq compiler internally makes transformations of the following form:

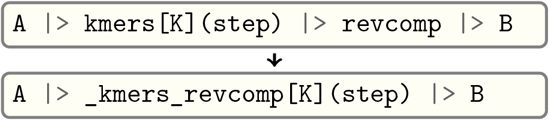

where `_kmers_revcomp[K]` is a function that iterates over reverse complemented *k*-mers by first reverse complementing the entire sequence *then k*-merizing.

The effect of these optimizations is shown in Figure 5d, which shows the performance of a *k*-merization and reverse complementation pipeline for a naive implementation (i.e. every *k*-mer is encoded and reverse complemented individually), an implementation using Algorithm 4 for *k*-merization, and finally Seq’s implementation, which uses both Algorithm 4 and the reverse complement optimization described above.

### Canonical k-mers

A “canonical” *k*-mer is the minimum (often, lexicographic minimum) of a *k*-mer and its reverse complement. Canonical *k*-mers are frequently used in *k*-mer counting algorithms as well as various hashing schemes to ensure that *k*-mers are treated as equal to their reverse complements [24, 38]. For example, many *k*-mer counting methods report counts of canonical *k*-mers rather than raw *k*-mer counts. Consequently, a common pipeline pattern is to pipe the output of `kmers` to the `canonical` function, which simply returns the canonicalization of its argument.

A naive implementation of such a pipeline might reverse complement each *k*-mer individually, but a better approach is to use two sliding windows when iterating over *k*-mers: one for the forward-direction *k*-mer, and one for the reverse complement. When the step size is 1, for instance, the next base is shifted in on the right of the forward window, while the *complement* of the base is shifted in on the *left* of the reverse window, thereby ensuring the reverse window always holds the reverse complement of the forward window. Using this approach, no *k*-mer ever actually needs to be reverse complemented during the iteration. The Seq compiler recognizes this pattern and makes use of a double-sliding-window implementation, `_kmers_canonical`:

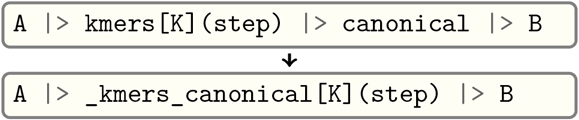

### Software prefetching

Large genomic indices, coupled with cache-unfriendly access patterns, often result in a substantial fraction of stalled memory-bound cycles in many genomics applications [33, 3, 40]. As a result, the Seq compiler performs optimizations to overlap the cache miss latency of an index lookup with other useful work, similar to the general-purpose approaches described in [17] and [12].

To illustrate the effect of this optimization, consider the pipeline `A |> B |> C` where function `B` performs an index lookup. In Seq, by using the @prefetch annotation on `B`, the function is implicitly transformed into a coroutine which performs a software prefetch, yields, then commences with the actual index lookup once it is resumed. Code for the pipeline itself is generated to include a dynamic scheduler, which manages multiple instances of the `B` coroutine, suspending and resuming them as needed until all are complete. In this way, while one instance of `B` is performing its prefetch, another can continue doing useful work, compared to the original pipeline where progress would be stalled until a given index lookup completes. A concrete example for FM-indices is shown in Figure 7, wherein the typical backwards-extension FM-index search algorithm is applied to count occurrences of subsequences from an input FASTQ file.

The performance of the code from Figure 7 for various values of *k*, both with and without the @prefetch line, is shown in Figure 5e. For smaller *k*, the “head” of the FM-index is largely cache-resident, so the prefetch optimization has no benefit. For larger *k*, however, the effect of cache misses increases and the optimization significantly reduces runtime (by almost 2× at *k*= 32). The prefetch optimization also works with other types of indices, such as hash tables, although we found it to be most effective on tree-like structures like FM-indices.

### Inter-sequence alignment

Sequence alignment via a dynamic programming algorithm like Smith-Waterman is arguably one of the most prevalent operations in genomics applications, and has been the subject of much research and optimization, particularly with regards to SIMD parallelization [36, 13, 32]. There are two approaches to SIMD alignment, referred to as *intra-sequence* and *inter-sequence*. An intra-sequence implementation uses SIMD to align a single pair of sequences, whereas an inter-sequence approach aligns a batch of pairs at once, and uses SIMD across different sequence pairs rather than within one pair. Inter-sequence alignment can be substantially faster than intra-sequence, but is rarely used in practice due to programming difficulty [37].

In Seq, however, inter-sequence alignment is just as easy to program as intra-sequence, as shown in Figure 7. When a function marked @inter_align is used in a pipeline, the Seq compiler will perform transformations similar to those used for software prefetching; specifically, the function is transformed into a coroutine and suspended just before the alignment. During execution of the pipeline, several hundred suspended coroutines are batched, have their sequences aligned using inter-sequence alignment, and are resumed after the alignment result is returned to them. Similar to the prefetch optimization, this is orchestrated by a dynamic scheduler of coroutines, the code for which is generated in the body of the pipeline. An illustration of such a pipeline before and after this transformation is shown in Figure 7 (bottom right).

Performance of inter-sequence alignment compared to intra-sequence is shown in Figure 5f, where inter-sequence is shown to be almost 4× faster in some cases. The code difference between the two implementations used in this figure is just a single line: the @inter_align annotation on the function performing the alignment. Note that Seq uses KSW2 [23, 36] as its intra-sequence alignment kernel.

This approach to alignment also lends itself well to other backends. For example, it would be possible to instead perform inter-sequence alignment on a GPU or FPGA rather than a CPU, and tune the parameters (e.g. sequence buffer size) for each backend. A domain-specific language and compiler such as Seq makes it possible to explore various backends and their benefits in a systematic way, and we plan to pursue this as future work.

## Performance benchmarking

To evaluate runtime and memory usage of our Seq implementations, all experiments were run on a dual-socket system with Intel Xeon E5-2695 v2 CPUs (2.40 GHz) with 12 cores each (totalling 24 cores and 48 hyper-threads) and 377GB DDR3-1066 RAM with 30MB LLC per socket. The Seq version is 0.9.7.

## Code availability

Seq is freely available at https://seq-lang.org. We provide the source code for each of the Seq implementations presented here on GitHub at https://github.com/seq-lang/seq/tree/develop/test/apps, with the exception of HapTree-X, which is available at https://github.com/0xTCG/haptreex.

## Acknowledgments

The authors thank Deniz Yorukoglu for helpful discussions, as well as Brian Hie for invaluable comments and edits on the main text.

## Correspondence

Correspondence and requests for materials should be addressed to Saman Amarasinghe and Bonnie Berger.

## Results

### *k*-merization Algorithm

**Algorithm 4:**
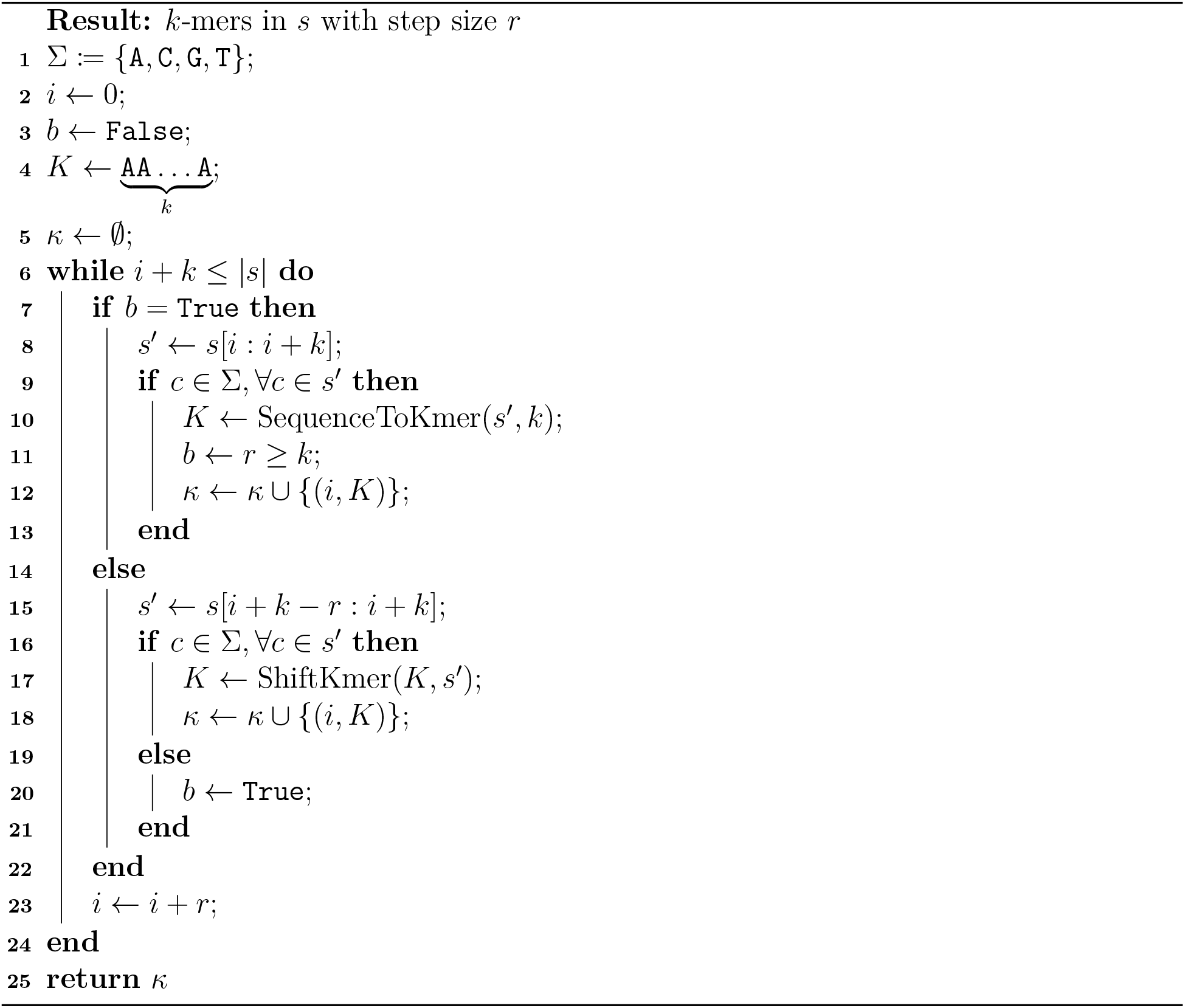
*k*-mers contained in sequence *s* at offsets a multiple of *r*, skipping non-`ACGT` bases. For simplicity, we ignore reverse complementation and assume *s* is in the forward direction.

## Footnotes

2 We found experimentally that sequences longer than 512 bases did not benefit from inter-sequence alignment. Note that this limit is user-adjustable.

